# Functional Anatomy of the TDP-43 Redox Sensor

**DOI:** 10.1101/2021.12.05.471332

**Authors:** Xiaoming Zhou, Lily Sumrow, Lillian Sutherland, Daifei Liu, Tian Qin, Steven L. McKnight, Glen Liszczak

**Affiliations:** Department of Biochemistry, UT southwestern Medical Center, Dallas, TX 75390, USA

## Abstract

TAR binding protein 43 (TDP-43) is an RNA binding protein that assists in the maturation, export and sub-cellular localization of mRNA. The carboxyl terminal 153 residues of TDP-43 are of low sequence complexity and allow for self-association of the protein in a manner leading to its phase separation from an aqueous environment. These interactions assist TDP-43 in forming cytoplasmic RNA granules involved in the transport of mRNA for localized translation. Self-association of the TDP-43 low complexity (LC) domain is facilitated by a region of twenty five residues that are of extreme evolutionary conservation. The molecular basis for self-adherence of the protein through this region has been illuminated by a combination of structural and biochemical studies, allowing definition of a morphologically specific cross-β structure predicted to be weakly assembled by main chain hydrogen bonds. In this study we have investigated the importance of individual, Pauling hydrogen bonds hypothesized to facilitate self-adherence of the TDP-43 LC domain.

## Introduction

Messenger RNA (mRNA) molecules made in the nucleus of eukaryotic cells must be exported to the cytoplasm to enable translation. A growing list of mRNAs have been found to be translated in a localized manner, such that newly formed proteins are produced in locations proximal to their intended sites of need (*1*). During the process of movement from nuclear synthesis to eventual translation, mRNAs can be transiently housed in granular particles visible by light microscopy (*1*). These RNA granules are not encased within lipid membranes, and considerable interest is being devoted to the challenge of understanding how they are organized.

The prominence of TDP-43 as an mRNA binding protein is reflective of two facts. First, it has long been recognized as a component of neuronal RNA granules that move mRNA molecules along the dendrites of neurons to facilitate enhanced translation at active synapses (*2-5*). Second, TDP-43 is frequently found in an aggregated state in the brain tissue of patients suffering from neurodegenerative disease (*6, 7*).

A carboxyl terminal region of the TDP-43 protein has been implicated in both its partitioning into RNA granules and its aberrant aggregation as a function of disease. The carboxyl terminal 153 residues of TDP-43 are of low sequence complexity and were long thought to be intrinsically disordered (*8*). It is this region of the protein that is able to self-associate in a manner driving phase separation. The relationship of the TDP-43 low complexity (LC) domain to pathologic aggregation has been supported by the discovery of missense mutations within it causative of neurodegenerative disease (*9, 10*).

That the TDP-43 LC domain may help facilitate RNA granule formation is evidenced by its ability to become phase separated from aqueous solution. Upon incubation at high concentration under conditions of neutral pH and physiologically relevant monovalent salts, the TDP-43 LC domain quickly forms liquid-like droplets that mature into a gel-like state (*11, 12*). This behavior is reminiscent of the LC domains associated with many other RNA binding proteins that have likewise been implicated in the organization of RNA granules (*13-15*).

The LC domain of TDP-43 is different in one conspicuous regard from prototypic LC domains that phase separate. The latter proteins are routinely enriched with tyrosine and/or phenylalanine residues functionally important for phase separation (*13, 16-19*). Instead of being enriched in aromatic amino acids, the LC domain of TDP-43 contains ten evolutionarily conserved methionine residues. This feature imparts oxidation sensitivity to phase separated liquid-like droplets and hydrogels formed from the TDP-43 LC domain (*11*). When exposed to hydrogen peroxide (H_2_O_2_), TDP-43 droplets melt in a manner correlated to the formation of methionine sulfoxide adducts. Upon chemical reduction by methionine sulfoxide reductase enzymes, thioredoxin, thioredoxin reductase and NADPH, the TDP-43 LC domain again forms phase separated droplets. It has been hypothesized that this form of post-translational modification of the TDP-43 LC domain may define a redox switch allowing for locally controlled dissolution of RNA granules (*11*).

A segment of twenty five amino acids defines the region of the TDP-43 LC domain critical for self-association and phase separation. This region has been perfectly conserved through the 400M years of evolutionary divergence between fish and humans. This region has also been observed by chemical footprinting to assume a structurally ordered state in phase separated liquid-like droplets, hydrogels and living cells (*11*). Precisely the same region of the TDP-43 LC domain has been reported to form the molecular core of a morphologically specific cross-β structure (*20*).

These concordant experimental observations point to the utility of a small and simple molecular device useful for specific aspects of cellular organization and biological regulation. The molecular assembly enabling self-association of the TDP-43 LC domain, like all cross-β structures, should be reliant on intermolecular hydrogen bonds adhering adjacent β strands. Here we report the results of experiments designed to test this assumption.

## Results

A purified protein fragment corresponding to the intact low complexity (LC) domain of TDP-43 forms phase separated liquid-like droplets within seconds after suspension at room temperature in a neutral pH buffer supplemented with 150 mM NaCl. As compared with denatured, unfolded protein, methionine residues 322 and 323 are protected from H_2_O_2_-mediated oxidation in phase separated preparations of the TDP-43 LC domain (*11*). These oxidation-resistant methionine residues co-localize with a dagger-like cross-β structure composed of 16 residues observed in three cryo-EM reconstructions of polymers formed from the TDP-43 LC domain (*20*). The dagger-like structure and locus of oxidation protection themselves co-localize to a region of 25 residues corresponding to the most evolutionarily conserved region of the TDP-43 LC domain.

Similar patterns of oxidation protection of methionine residues 322 and 323 of the TDP-43 LC domain have been observed in liquid-like droplets, labile cross-β polymers and living cells (*11*). Recognizing co-localization of these two methionine residues within the three structures resolved by cryo-EM, we hypothesized that self-association and consequential phase separation of the TDP-43 LC domain might be driven by an assembly defined by the small cross-β structure itself.

In efforts to experimentally investigate this hypothesis, we prepared twenty three variants of the TDP-43 LC domain differing only in chemical modification of the peptide backbone nitrogen atom of each residue housed within the ultra-conserved region of the protein. Variants carried a methyl cap on individual peptide backbone nitrogen atoms, thus allowing for a systematic analysis of the involvement of Pauling hydrogen bonding for self-association and phase separation of the protein into liquid-like droplets. It is obvious from cryo-EM images of TDP-43 LC domain polymers that inter-peptide hydrogen bonds represent favorable adhesive forces holding protein chains together in a parallel, in-register cross-β conformation. Our experimental strategy allowed assessment as to whether hydrogen bonds located either within or outside of the cross-β structure might be required for phase separation.

Variants of the TDP-43 LC domain capped at individual backbone nitrogen atoms were assembled using a three-piece native chemical ligation strategy. Briefly, synthetic peptide thioesters bearing single N^α^-methyl amino acid variants were ligated to N- and C-terminal flanking fragments of the TDP-43 LC domain that had been expressed in bacterial cells (see legend to Fig. 1). Ligation products were purified by HPLC and analyzed by mass spectrometry to confirm the addition of a single backbone methylation site.

**Fig. 1.**
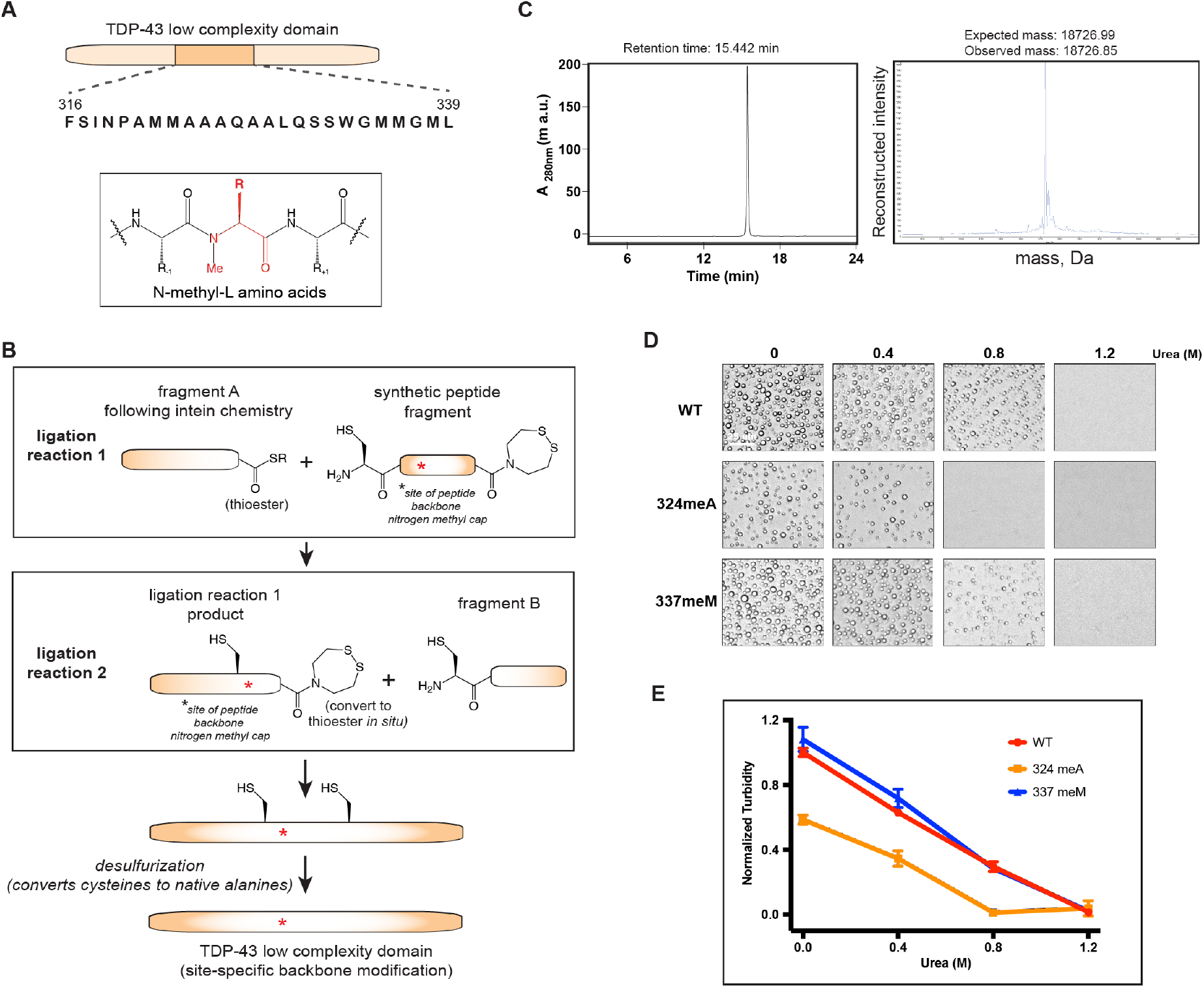
Preparation of semi-synthetic derivatives of the TDP-43 LC domain carrying single, methyl-capped peptide backbone nitrogen atoms. The LC domain of TDP-43 is encompassed by the C-terminal 153 residues of the protein, extending from residue 262 through 414 **(Panel A)**. N^β^-methyl amino acid scanning analysis focused on an ultra-conserved sequence of the LC domain extending from residue 316-339, the amino acid sequence of which is shown in **Panel A**. N^β^-methyl amino acids (red color and red asterisks) were introduced into synthetic peptides that were conjugated via sequential native chemical ligation reactions to flanking protein fragments produced in bacteria **(Panel B)**. Detailed methods describing protein semi-synthesis, purification and mass spectrometry are described in Materials and Methods. Ligation products contained cysteine residues in place of alanine residues at both ligation boundaries. Both cysteine residues were chemically desulfurized to alanine such that the only difference between engineered LC domains and the native LC domain was the presence of a single N^β^-methyl group at the desired site. Ligation products were purified by HPLC allowing confirmation of chemical composition by high resolution, intact mass spectrometry **(Panel C)**. Light microscopic analyses of phase separation for three representative samples are shown in **Panel D**, including the reconstructed, semi-synthetic, native TDP-43 LC domain, a variant carrying a methyl cap on the peptide backbone nitrogen atom of alanine residue 324, and a variant carrying a methyl cap on the peptide backbone nitrogen of methionine residue 337. Each protein was incubated under conditions of neutral pH and physiologically normal monovalent salts allowing for formation of phase separated liquid-like droplets. Droplet formation was assayed in triplicate in normal buffer as well as buffers supplemented by 0.4 M urea, 0.8 M urea and 1.2 M urea. Quantitative analysis of phase separation for all protein samples under the four experimental conditions was monitored by spectroscopic analysis of turbidity **(Panel E)**. Scale bar = 25 µm.

Following purification, samples were assayed for the formation of phase separated liquid-like droplets in the presence of a standard buffer as well as buffers supplemented with 0.4 M, 0.8 M or 1.2 M urea. As shown in Fig. 1, the intact LC domain of TDP-43 formed liquid-like droplets in native buffer. Fewer droplets were observed at 0.4 M urea, fewer still at 0.8 M urea, and no liquid-like droplets were observed in buffer supplemented with 1.2 M urea. Droplet formation was quantified by a simple spectroscopic assay of turbidity.

Compared with the native LC domain of TDP-43, the variant bearing a methyl cap on the peptide backbone nitrogen associated with alanine residue 324 formed fewer droplets in native buffer, fewer droplets in buffer supplemented with 0.4 M urea, and no droplets in buffer supplemented with either 0.8 M or 1.2 M urea. Quantitation of droplet formation by spectroscopic measurements of turbidity confirmed that the variant bearing a methyl cap on the peptide backbone nitrogen associated with alanine 324 was diminished in its capacity to phase separate under the three conditions supportive of droplet formation by the intact TDP-43 LC domain. By contrast, a variant of the TDP-43 LC domain bearing a methyl cap on the peptide backbone nitrogen associated with methionine residue 337 formed morphologically indistinguishable droplets as compared with the native protein under all conditions tested. Droplet numbers quantified by turbidity for the 337meM variant were also indistinguishable from the native protein.

Twenty three variants of the TDP-43 LC domain bearing single methyl caps spanning residues 316-339 of the TDP-43 LC domain were prepared via our three-piece assembly strategy and assayed for phase separation. This twenty four residue region corresponds to the most evolutionarily conserved segment of the TDP-43 LC domain. The only amino acid not evaluated in this manner was proline residue 320. Proline, which has a side chain that is twice connected to the peptide backbone to form a pyrrolidine ring, is the only amino acid devoid of the NH group required for Pauling hydrogen bonding.

The native TDP-43 LC domain sample and all twenty three variants were evaluated for phase separation in triplicate, allowing error bars to validate assay reproducibility. Microscopic images of all proteins as observed in normal buffer and the three different concentrations of urea are shown in Supplemental Data, Fig. S1. The turbidity assays gathered in triplicate for all twenty four protein samples are shown in Fig. 2. These data reveal nine variants as being significantly impaired in their ability to phase separate, two variants as being partially impaired, and twelve variants as phase separating in a manner similar to the native protein.

**Fig. 2.**
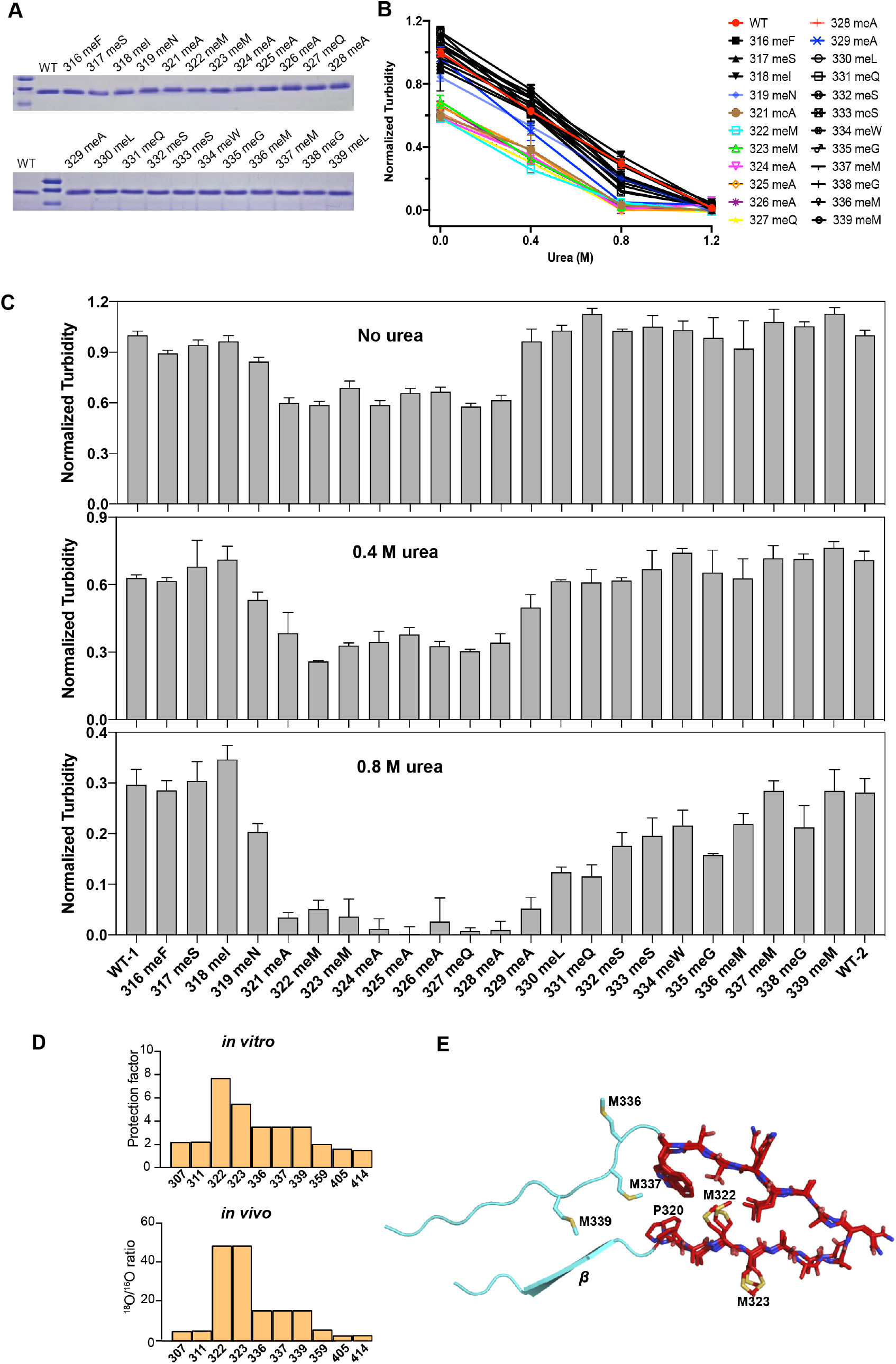
Phase separation capacities of twenty three variants of the TDP-43 low complexity domain differing according to the presence of a single, methyl-capped peptide backbone nitrogen atom. Twenty three variants of the TDP-43 LC domain were compared with the native protein in assays of phase separation. All proteins were evaluated by SDS-PAGE to confirm protein purity and integrity **(Panel A)**. Five minutes after suspension in buffer of neutral pH and physiological monovalent salt, samples were spectroscopically measured for turbidity **(Panel B)**. In addition to standard buffer, each sample was further analyzed in buffers supplemented by 0.4 M urea, 0.8 M urea and 1.2 M urea. All samples were analyzed in triplicate and further evaluated and photographed by light microscopy (Supplemental Data, Fig. S1). Turbidity measurements revealed two classes of variants irrespective of the presence or absence of urea. One class composed of nine variants showed, relative to the native TDP-43 LC domain, reduced turbidity and fewer liquid-like droplets under all conditions. The other class, composed of thirteen variants, showed evidence of phase separation indistinguishable from the native protein. The histograms shown in **Panel C** reveal the normalized, average turbidity measurements of twenty four protein samples as evaluated in the absence of urea as well as in the presence of either 0.4 M or 0.8 M urea. Turbidity is graphed on the Y axis of each plot relative to the amino acid sequence of the ultra-conserved region of the TDP-43 LC domain between phenylalanine residue 316 and methionine residue 339 (X axis). Irrespective of assay condition, all nine of variants exhibiting impediments to phase separation clustered between alanine residue 321 and alanine residue 329. **Panel D** shows reproductions of published data pinpointing the region of the TDP-43 LC domain footprinted to reveal structure-dependent protection from methionine oxidation (*11*). Structure-mediated oxidation protection, as observed both *in vitro* and in living cells, extends from methionine residue 322 through methionine residue 339 of the TDP-43 LC domain. **Panel E** shows the molecular structure of a cross-β polymer formed from the TDP-43 LC domain as resolved by cryo-electron microscopy (*20*). The structurally ordered region of the protein extends from proline residue 320 to methionine residue 337.

The histograms of Fig. 2, Panel C show normalized turbidity measurements for the twenty four protein samples (Y axis) relative to residue positions along the protein chain (X axis). One histogram shows normalized turbidity data collected in the absence of urea, one shows data collected in the presence of 0.4 M urea, and one shows data collected in the presence of 0.8 M urea. All three histograms reveal that all nine of the significantly impaired variants cluster between alanine residue 321 and alanine residue 329. Panels D and E of Fig. 2 further reveal that the region wherein removal of single hydrogen bonds impedes phase separation co-localizes with the footprinted region of structure-dependent oxidation protection, and the Eisenberg cross-β structure resolved by cryo-EM (*11, 20*).

The new experimental observations reported in this study give direct evidence of the importance of main chain hydrogen bonding for self-association and phase separation of the TDP-43 LC domain. We are aware of no other studies that have specifically addressed the importance of Pauling hydrogen bonding to LC domain phase separation. That elimination of single hydrogen bonds can reproducibly yield distinct phenotypic deficits gives evidence of the inherent lability of the cross-β structure responsible for self-association of the TDP-43 LC domain.

Whereas data presented in Figures 1 and 2 give evidence that Pauling hydrogen bonding is indeed important for self-association of the TDP-43 LC domain, they fail to address the potential importance of amino acid side chains. In the absence of side chain-mediated chemical interactions, there would be no opportunity for specificity in formation of the cross-β structure itself. In efforts to investigate the functional involvement of amino acid side chains, we performed glycine scanning mutagenesis across the ultra-conserved region of the TDP-43 LC domain. Among the twenty amino acids that might have been used for scanning mutagenesis, we chose glycine because of the chemical simplicity of its side chain.

The pattern of effects of twenty six side chain variants of the TDP-43 LC domain upon phase separation was similar to the pattern of effects resulting from hydrogen bond denial (Fig. 3). Substitution of the native residues between alanine 324 and leucine 330 by glycine led to the formation of distinctly misshapen droplets. It was this same region of the protein that was most sensitive to methyl capping of individual peptide backbone nitrogen atoms (Fig. 2).

**Fig. 3.**
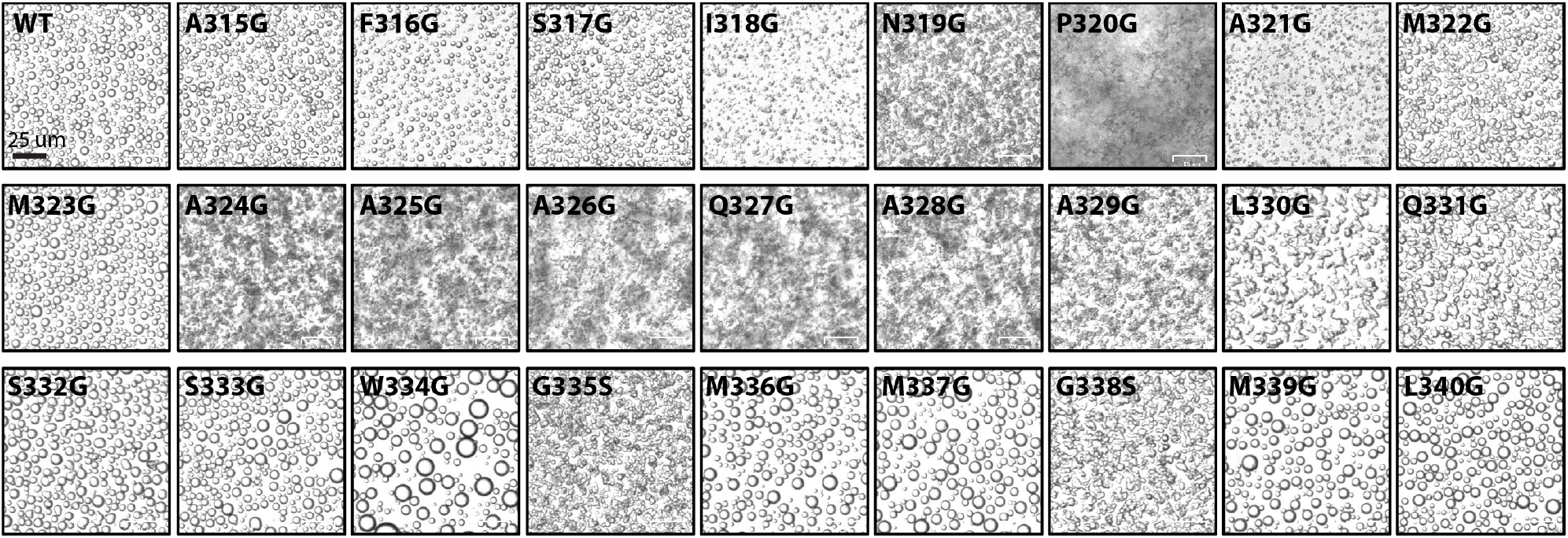
Phase separation capacities of twenty six variants of the TDP-43 LC domain differing according to the replacement of a single amino acid residue with glycine. Single amino acid residues within the ultra-conserved region of the TDP-43 LC domain were changed to glycine by conventional mutagenesis and expression in bacterial cells. Two of the twenty six positions already contained glycine (G335 and G338) in the native sequence and were mutationally changed to serine. Each protein was purified and tested for phase separation in buffer of neutral pH and physiologically normal monovalent salt ions. Light microscopic images were photographed 1 h subsequent to initiation of each reaction. In addition to assays performed in standard buffer, each protein sample was also evaluated for its capacity to phase separate in the presence of 0.4 M urea, 0.8 M urea and 1.2 M urea (Supplemental Data, Fig. S2). Scale bar = 25 µm.

Of the ten variants located on the carboxyl-terminal side of leucine 330, only two caused notable changes in droplet morphology (G335S and G338S). Perhaps coincidentally, or perhaps importantly, these are the only positions represented by glycine in the native sequence of TDP-43 (and were changed to serine as part of the mutagenesis scan).

Of the nine variants located on the amino-terminal side of alanine 324, only two caused notable changes in phase separation. Both the N319G and P320G variants led to the formation of misshapen droplets. Among all twenty six variants tested, the P320G variant revealed the most profound effect on droplet formation. Immediately following suspension in aqueous buffer, the P320G variant formed a heavily tangled precipitate.

In addition to inspecting the twenty six glycine scanning variants in normal aqueous buffer, we further analyzed droplet morphology in buffers supplemented with 0.4 M, 0.8 M and 1.2 M urea. As shown in Supplemental Data Fig. S2, the pattern of effects of glycine scanning variants was largely similar irrespective of the addition of urea. These added assays revealed that certain variants were, relative to the native protein, either more or less sensitive to the urea denaturant. In particular, the four “outlying” variants – N319G, P320G, G335S and G338S - continued to display either precipitates or misshapen droplets in the presence of 1.2 M urea (a condition that fully melts liquid-like droplets made from the native TDP-43 LC domain). Paradoxically, we also observed urea-resistant droplets for the S332G and S333G variants. In summary, these experiments confirm the importance of numerous amino acid side chains within the ultra-conserved region of the TDP-43 LC domain for the formation of labile and morphologically spherical, phase separated droplets.

Having observed profound deficits in phase separation assays for the P320G variant, we wondered whether mutation of other proline residues within the TDP-43 LC domain might also lead to significant deficits. In addition to P320, the sequence of the LC domain specifies proline residues at positions 280, 349 and 363 (Fig. 4B). Each of these residues was individually changed to glycine followed by expression, purification and assay of phase separation. None of the three variants caused any effect on either the morphology or stability of phase separated liquid-like droplets (Fig. 4C). We thus conclude that proline residue 320 is of unique importance to function of the TDP-43 LC domain.

**Fig. 4.**
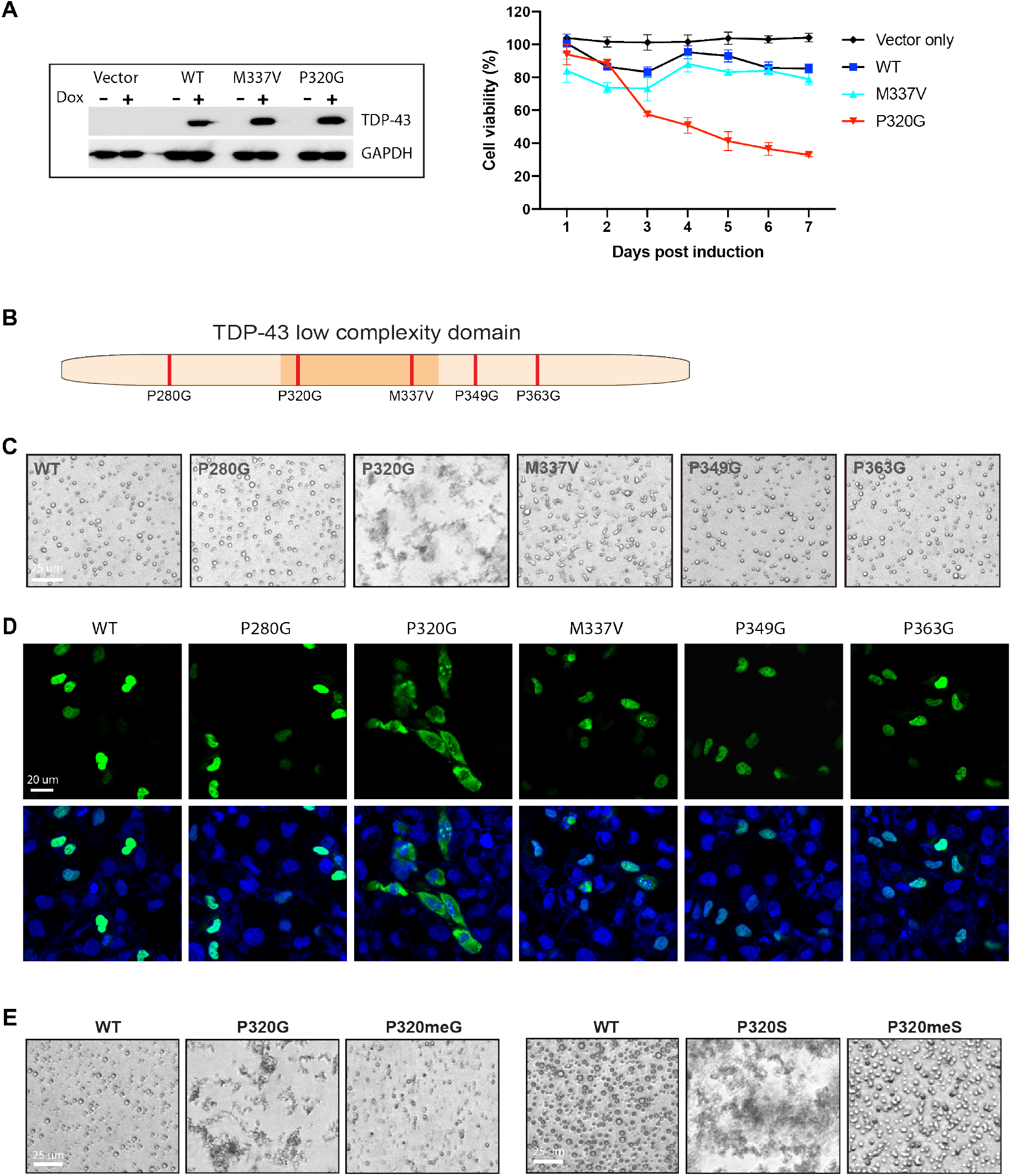
Evidence of the unusual importance of proline residue 320 of the TDP-43 LC domain. Cultured U2OS cells were stably transformed with vectors allowing for conditional, doxycycline-mediated expression of FLAG-tagged versions of native, full length TDP-43 (WT), the ALS-causing M337V variant of TDP-43, or the P320G variant causative of severe precipitation. Single transformant clones were isolated in the absence of doxycycline and screened for the purpose of finding stable clones that expressed equivalent protein levels post-doxycycline induction (Materials and Methods). Cell lysates were recovered before and twenty four hours after doxycycline induction, run on an SDS-PAGE gel, and Western-blotted using a FLAG antibody or antibodies to GADPH as a loading control **(left side, Panel A)**. Cell viability was monitored daily post-induction for each clone, revealing minimal effects from doxycycline-induced expression of either the native (WT) TDP-43 protein or the ALS-causing M33V variant, as compared with substantial deficits resulting from expression of the P320G variant **(right side, Panel A)**. The native TDP-43 protein contains proline residues at three other positions within its LC domain, all located outside of the ultra-conserved domain shaded in amber **(Panel B)**. Proline-to-glycine variants at residue 280, 349 or 363 exhibited no deficits in phase separation **(Panel C)**. Constitutive expression vectors encoding fusion proteins linking GFP to the amino terminus of full-length TDP-43 were transiently transfected into U2OS cells. Nucleus-restricted GFP was observed for native TDP-43 linked to GFP, as well as for the proline-to-glycine variants of residues 280, 349 and 363, and the ALS-causing M337V variant. Transient expression of the P320G variant led to predominantly cytoplasmic GFP staining, with residual nuclear staining appearing aberrantly punctate **(Panel D)**. Semi-synthesis of variants of the TDP-43 LC domain bearing either proline-to-glycine or proline-to-serine changes at residue 280 formed tangled precipitates upon tests of phase separation. Methyl capping of the peptide backbone nitrogen atom associated with the glycine residue of the P320G variant, or the peptide backbone nitrogen atom associated with the serine residue of the P320S variant, yielded proteins for which formation of phase separated liquid-like droplets was restored **(Panel E)**. Scale bar in Panel C = 25 µm, scale bar in panel D = 20 µm, and scale bar in panel E = 25 µm.

A number of well-validated, amyotrophic lateral sclerosis (ALS)-causing missense mutations map within or close to the ultra-conserved region of the TDP-43 LC domain (*9*). Included among such mutations is the particularly well-studied M337V variant that leads to the formation of oxidation-resistant liquid-like droplets (*11*). These observations raise concern as to why human genetic studies have failed to report ALS-causing missense mutations in proline residue 320?

Given its severe propensity to aggregate into stable precipitates, we considered the possibility that expression of the P320G variant might be incompatible with cell viability. To this end U2OS cells were stably transformed with inducible transgenes encoding native TDP-43, the M337V ALS-causing variant, or the P320G variant. After isolating cell clones expressing comparable levels of the three proteins upon conditional induction, and no detectable expression of the proteins in the absence of the chemical inducer of expression, we performed viability studies as a function of time post-induction. As shown in Fig. 4A, conditionally induced expression the P320G variant substantially reduced cell viability. No such impediment to viability was observed for cells transformed with the parental expression vector, nor for cells conditionally expressing either the native TDP-43 protein or the ALS-causing M337V variant. We simplistically conclude that the P320G variant has not appeared from human genetic studies as an ALS-causing missense mutation for the simple reason that this form of the protein is cell-lethal.

To investigate the behavior of the P320G variant in living cells, as compared with the native protein, the ALS-disposing M337V variant, and the three P-to-G variants at positions 280, 349 and 363, expression vectors were prepared linking each variant of TDP-43 to green fluorescent protein (GFP). U2OS cells were transiently transfected with each expression vector. Twenty four hours later the cells were visualized by confocal microscopy as shown in Fig. 4D. GFP signal associated with the fusion protein linked to the native TDP-43 protein was restricted to the nuclei of transfected cells. These methods also revealed nuclear staining for the ALS-causing M337V variant. By contrast, the GFP signal for the P320G variant was almost exclusively restricted to the cytoplasm, with residual nuclear staining confined to prominent nuclear puncta. Finally, the P280G, P349G and P363G variants yielded patterns of nuclear staining indistinguishable from the native TDP-43 protein.

Why is proline residue 320 of such unusual importance to function of the TDP-43 LC domain? Recognizing that proline is the only amino acid not permissive of forming Pauling hydrogen bonds, its replacement by any other amino acid would obviously restore hydrogen bonding capacity. In addition to the P320G variant characterized thus far, we also mutated proline 320 to either serine or alanine. Both variants caused equally profound precipitation of the TDP-43 LC domain relative to the P320G variant in phase separation assays (Supplemental Data, Fig. S3). These observations raised the possibility that proline residue 320 might function as a chemical insulator functioning to locally terminate one end of the dagger-like cross-β structure.

If the glycine or serine substitutions of proline residue 320 exert their deleterious effects by regenerating hydrogen bonding capacity, this deficit should be overcome were we to methylate the peptide backbone nitrogen atom of the glycine residue in P320G, or the serine residue in P320S. We again employed the three-piece TDP-43 ligation reaction detailed in Fig. 1 to introduce a methyl cap onto the peptide backbone nitrogen atom of these two variant residues. These efforts led to the rescue of phase separated liquid-like droplets in both cases (Fig. 4E).

## Discussion

By devising and deploying methods of semi-synthetic protein chemistry, we have systematically evaluated the importance of twenty three peptide backbone amide protons within the TDP-43 LC domain for formation of phase-separated liquid-like droplets. Denial of hydrogen bonding capacity in a localized region of the TDP-43 LC domain distinctly weakened protein self-association. This region of sensitivity to the focused removal of hydrogen bonds falls within a known cross-β structure. As such, our observations comport with the proposed involvement of the small cross-β structure in self-association and phase separation by the TDP-43 LC domain. These observations further conform to previous studies of the fused-in-sarcoma (FUS) and hnRNPA2 LC domains that also employ labile cross-β structures accounting for the mechanistic basis of phase separation (*11, 15, 21*).

In addition to studies focused on Pauling hydrogen bonding, glycine scanning mutagenesis experiments were performed to assess the importance of amino acid side chains to the phenomena of LC domain self-association and phase separation. The results of such studies confirm the importance of many amino acid side chains, particularly those proximal to the region of the TDP-43 LC domain that is both protected from methionine oxidation in the self-associated state and responsible for formation of the cross-β structure.

These observations, even when combined with the results of our experiments deployed to systematically evaluate the importance of Pauling hydrogen bonds, do not constitute proof of the accuracy or biologic validity of the dagger structure. Further experimentation will be required to confidently believe that this structure properly and precisely defines how the TDP-43 LC domain self-associates. Despite this uncertainty, we reiterate four facts: (i) the region of the TDP-43 LC domain that is most strongly protected from H_2_O_2_-mediated oxidation extends from residue 322 through residue 339 (*11*); (ii) the region of the protein that forms the cross-β dagger structure extends from residue 320 through residue 337 (*20*); (iii) the region of the protein most sensitive to denial of hydrogen bonding capacity extends from residue 321 through residue 330 (Fig. 2); and (iv) the region of the protein most sensitive to systematic alteration of amino acid side chains extends from residue 319 through residue 338 (Fig. S2). In aggregate, these data cause us to strongly believe that self-association and phase separation of the TDP-43 LC domain is mediated by the formation of a labile cross-β structure located in immediate proximity to both the Eisenberg dagger and the region protected from methionine oxidation only when the LC domain is allowed to adopt its molecular structure.

Whereas the same region of the TDP-43 LC domain is sensitive to systematic elimination of the capacity for Pauling hydrogen bonding (Fig. 2 and Fig. S1), and systematic variation of side chains (Fig. S2), close inspection of the data reveals that the different methods of disruption led to different trends of effect. If a variant resulting from peptide backbone methylation had any phenotypic effect on phase separation, the effect universally weakened liquid-like droplets (Fig. 2 and Fig. S1). By contrast, the majority of side chain variants caused the formation of droplets that were more urea-resistant than the native TDP-43 LC domain.

A second difference between variants made by the different approaches was observed for droplet morphology. Denial of the capacity for Pauling hydrogen bonding tended to yield liquid-like droplets having spherical morphology. Almost all side chain variants yielded droplets having either misshapen morphology, or amorphous precipitates as exemplified by the P320G variant. We offer that spherically homogeneous droplet morphology may represent a surrogate assay for formation of the properly organized molecular assembly. Further experimentation will be required to either validate or refute this simplistic concept.

Quite by accident, we discovered the unusually extreme importance of proline residue 320. If this residue is changed to any other amino acid, the missing opportunity for Pauling hydrogen bonding is restored. This, we offer, represents the importance of proline 320. By lacking the backbone amide proton essential for Pauling hydrogen bonding, P320 is predicted to terminate the network of hydrogen bonds employed to form the dagger structure. Having noticed an unpaired β strand on the immediate, N-terminal side of P320 (Fig. 2E), we offer that restoration of the capacity for Pauling hydrogen bonding at this position might allow formation of a long and aberrantly stable cross-β structure. The properties of this aberrant structure are predicted to be sufficiently untoward to restrict TDP-43 to the cytoplasm and kill cultured mammalian cells.

That proline residue 320 of TDP-43 is of unusual importance to the function of its LC domain was emphasized, in a relative sense, by the results of mutational studies of three other proline residues located at positions 280, 349 and 363 within the LC domain. When any of these proline residues were changed to glycine, we observed no functional deficit in self-association/phase separation, and no discernable alteration of the intracellular localization of TDP-43 (Fig. 4). Our conclusion that proline residue 320 functions according to the absence of Pauling hydrogen bonding capacity is supported by methyl-capping of the glycine and serine residues of the P320G and P320S variants. In both cases, capping of these peptide backbone nitrogen atoms overcame otherwise aberrant precipitation, thus allowing formation of phase separated liquid-like droplets (Fig. 4E).

For the purpose of clarity, we make note of publications reporting that the TDP-43 LC domain can adopt an β-helical conformation in the same region we have shown to function via the formation of labile cross-β interactions (*8, 22, 23*). Evidence of β-helical conformation for the TDP-43 LC domain has been observed by solution NMR spectroscopy under conditions of acidic pH and in the complete absence of monovalent salt (deionized water). Two such studies have shown that when pH is raised to neutral, NMR spectra diagnostic of β-helical conformation disappear and are replaced by spectra diagnostic of β strand conformation(*8, 22*). We do not consider the conditions required for appearance of α-helical conformation within the TDP-43 LC domain to be of physiological relevance. The cytoplasm and nucleoplasm of living cells are of neutral pH, and consist of a mixture of monovalent and divalent salts in the range of 100-200 mM. All of our biochemical experiments have been performed under these physiologically relevant conditions, not at acidic pH in the complete absence of salt ions.

We close by emphasizing that we remain at an early stage in a quest to understand how the TDP-43 LC domain actually works. We are confident that it functions as a biological redox sensor, but are ignorant as to the cellular conditions wherein methionine oxidation would be employed to dissolve the self-associated state. We are likewise ignorant as to how, mechanistically, methionine oxidation causes melting of TDP-43’s labile, cross-β structure. The extreme evolutionary conservation the TDP-43 LC domain causes us to believe that we are studying a well-oiled molecular device that undoubtedly holds surprises to be unveiled in the years to come.

## Acknowledgments

We thank Masato Kato, Deepak Nijhawan and Rob Thompson for technical advice and encouragement. We thank Jugal Mohapatra and Kyuto Tashiro for assistance for synthesizing peptides. This work was supported by an anonymous donor and National Institute of General Medical Science of NIH grant R35 GM130358 and National Cancer Institute of NIH grant U54 CA231649 to SLM. This work was also supported by Welch Foundation grant I-2039-20200401 and I-2020-20190330 to GL and TQ, respectively, and a Cancer Prevention Research Institute of Texas grant RR180051 to GL.

## Data and materials availability

All data are available in the main text or the supplementary materials.

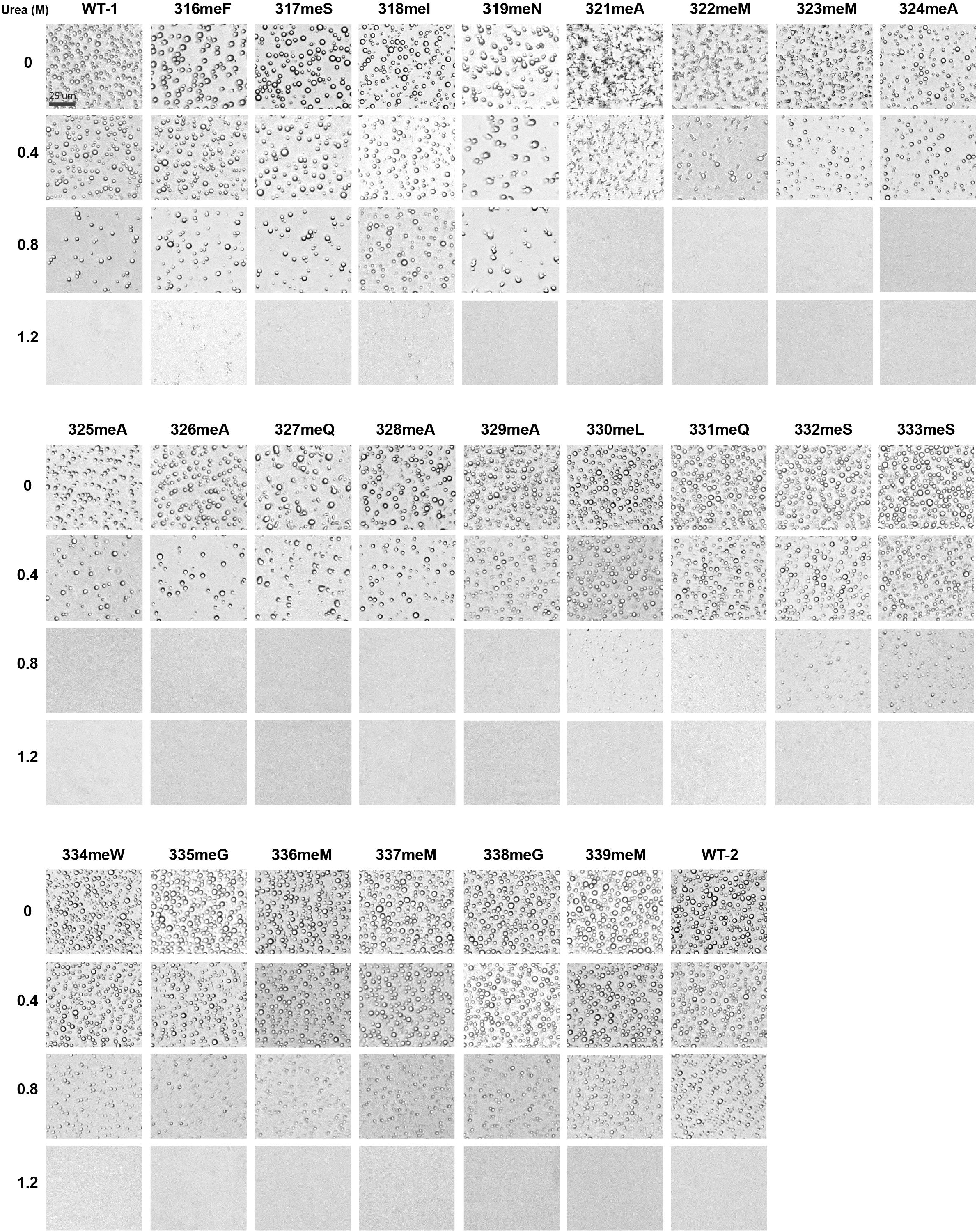

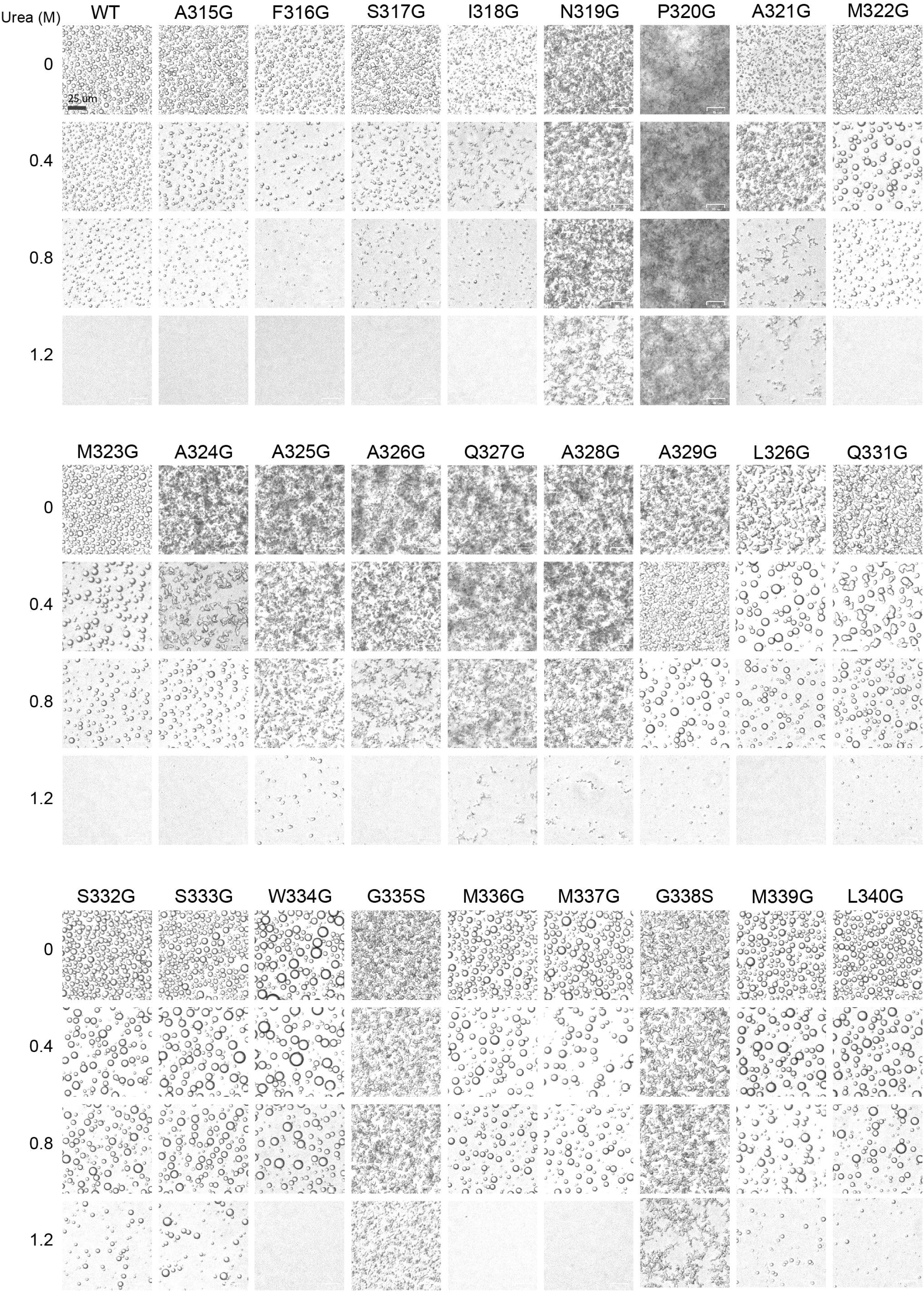

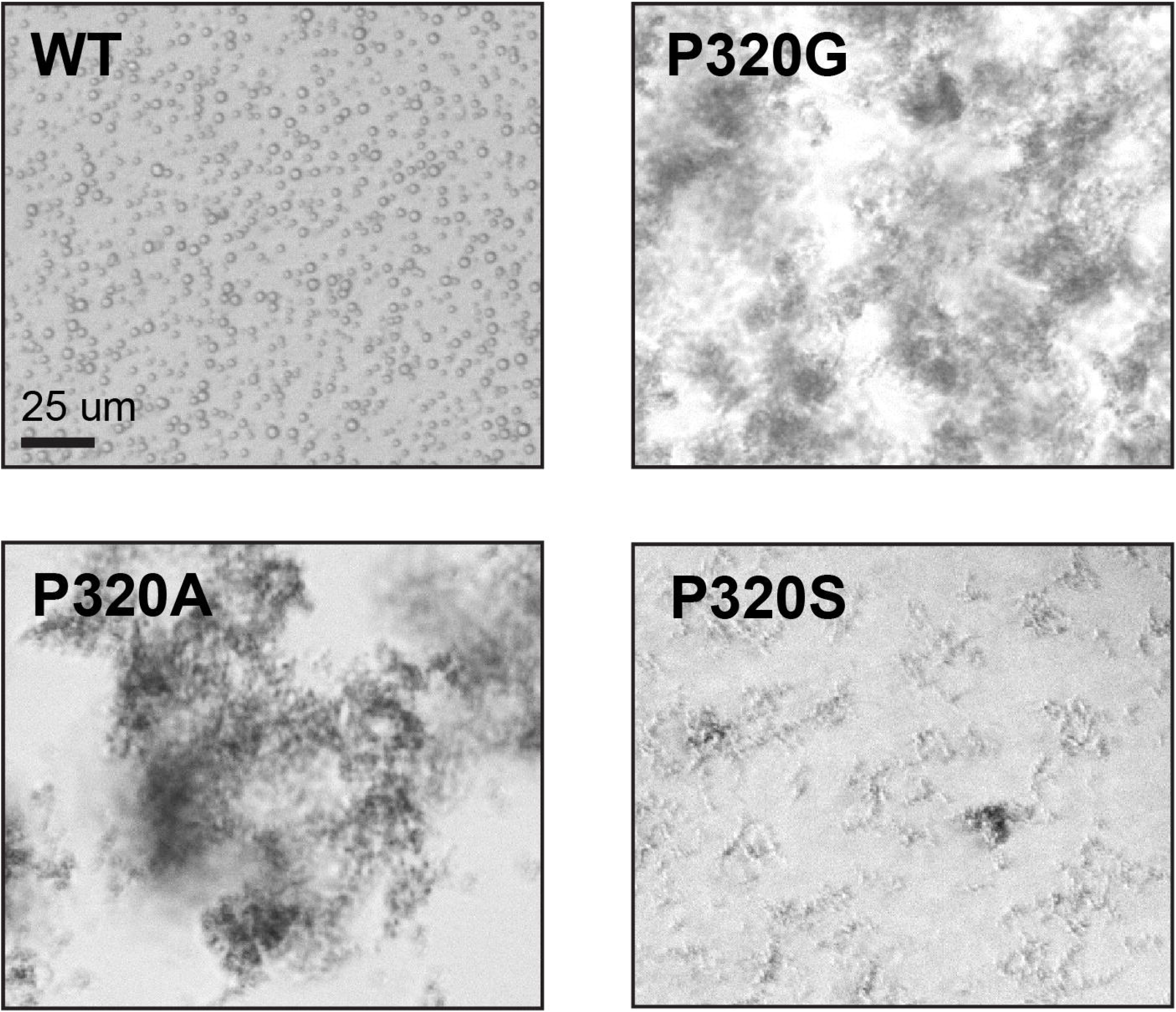

