## Supplementary material for "Functional Anatomy of the TDP-43 Redox Sensor": materials and methods, and supplementary figures

### Molecular cloning

Standard gene cloning and mutagenesis strategies were used to generate expression constructs for all recombinant proteins. For plasmid constructs used in the glycine mutagenesis scanning experiment, wild type TDP-43 low complexity domain (residues: 262-414) and its variants were cloned into pHis-parallel vector to yield His-TDP-43 low complexity domain (LCD) constructs. Recombinant GyrA intein fusion fragments of the TDP-43 low complexity domain that were required for expressed protein ligation were cloned into the pHis-parallel vector. The complete scan required two distinct three-piece ligation strategies and thus two separate intein fusion fragments (Strategy A, Piece 1, residues 262-314 and Strategy B, Piece 1, residues 262-326) of the TDP-43 LCD. The two separate, recombinant piece three constructs (Strategy A, Piece 3, residues 328-414 and Strategy B, Piece 3, residues 341-414) of the TDP-43 LCD were fused with a 6xHis-SUMO tag at N-terminus and a GryA intein at C-terminus in pET-28a vector. Wild type TDP-43 and mutant constructs for transient transfection of cultured U2OS cells were cloned as N-terminal GFP fusions in the pCDNA 3.1 vector with an N-terminal FLAG tag. For stable, inducible TDP-43 expression in mammalian cells, full Length WT, P320G, and M337V were constructed in the CMV mammalian expression vector containing the Tet operator and an N-terminal FLAG-GFP tag.

### Recombinant TDP-43 Piece 1 and Piece 3 sequences

Strategy 1, Piece 1: 6xHis-*TDP-43 (residues 262-314)*
SYYHHHHHHDYDIPTTENLYFQGAMDPEF*PKHNSNRQLERSGRFGGNPGGFGNQGGFG NSRGGGAGLGNNQGSNMGGGMNFG*-thioester

Strategy 2, Piece 1: 6xHis-*TDP-43 (residues 262-327)*
SYYHHHHHHDYDIPTTENLYFQGAMDPEF*PKHNSNRQLERSGRFGGNPGGFGNQGGFG NSRGGGAGLGNNQGSNMGGGMNFGAFSINPAMMAAAQA*-thioester

Strategy 1, Piece 3: *TDP-43 (residues 329-414; A329C)*
*CLQSSWGMMGMLASQQNQSGPSGNNQNQGNMQREPNQAFGSGNNSYSGSNSGAAIGWGSA SNAGSGSGFNGGFGSSMDSKSSGWGM*

Strategy 2, Piece 3: *TDP-43 (residues 341-414; A341C)*
*CSQQNQSGPSGNNQNQGNMQREPNQAFGSGNNSYSGSNSGAAIGWGSASNAGSGSGFNGG FGSSMDSKSSGWGM*

### Recombinant protein expression and purification

All recombinant proteins used in this study were expressed in *E.coli* BL21 (DE3) cells grown to an $OD_{600}$ of 0.6 in LB medium. Expression of His-TDP-43 LCD proteins and Piece 1 proteins were induced with presence of 0.8 mM IPTG at 37° C for 4 h. Expression of Piece 3 proteins was induced with 0.6 mM IPTG at 16° C for 16 h. The cells were harvested by centrifugation at 4,000 × g for 15 min.

For purification of His-TDP-43 LCD wild type and its variants, the cell pellets were resuspended in buffer A containing 25 mM Tris-HCl (pH 7.5), 200 mM NaCl, 6 M guanidine-HCl, 10 mM β-mercaptoethanol (β-ME), 20 mM imidazole, and disrupted by sonication. The cell lysates were clarified via centrifugation at 40,000 × g for 50 min. Supernatant was then applied to $Ni^{2+}$-NTA resin (Qiagen) and the column washed with buffer A and eluted with buffer A supplemented with 300 mM imidazole. Pure protein was concentrated by centrifugal filtration (Amicon Ultra-15) and samples were aliquoted, flash frozen, and stored at -80 °C for future use.

For purification of recombinant Piece 1 protein thioesters, the cell pellets were resuspended in buffer B, containing 25 mM potassium phosphate (pH 7.2), 150 mM NaCl, 8 M Urea, 1 mM TCEP, 20 mM imidazole, and disrupted by sonication. The resulting cell lysates were centrifugated at 40,000 × g for 50 min. The supernatants were applied to $Ni^{2+}$-NTA resin (Qiagen) and the column washed with buffer B and eluted with buffer B supplemented with 300 mM imidazole. The purified protein was refolded via overnight dialysis against a buffer containing 25 mM potassium phosphate (pH 7.2), 150 mM NaCl, 1 M Urea, 1 mM TCEP. The refolded protein was centrifugated at 4,000 × g for 30 min to remove precipitation, and the supernatant was supplemented with 300 mM MES-Na, 10 mM TCEP, adjusted to pH 7.0, and incubated at room temperature for 16 h to achieve thiolysis. Following thiolysis, protein solution was spun at 4,000 × g for 30 min to isolate precipitation. Precipitated protein was resolubilized in a buffer containing 25 mM potassium phosphate (pH 7.2), 150 mM NaCl, 6 M guanidine-HCl, 1 mM TCEP, and combined with the supernatant fraction and loaded onto $Ni^{2+}$-NTA resin. The column was washed with a buffer containing 25 mM potassium phosphate (pH 7.2), 150 mM NaCl, 6 M guanidine-HCl, 1 mM TCEP, 20 mM imidazole to remove unbound proteins. Piece 1-thioester was then eluted with the same buffer supplemented with 300 mM imidazole. The eluted Piece 1 fragment was supplemented with 1% TFA and purified by RP-HPLC (C4 preparative column). Fractions were analyzed by RP-HPLC and intact ESI-MS and pure fractions were lyophilized.

For purification of recombinant Piece 3 constructs, the cell pellets were resuspended in buffer C containing 25 mM potassium phosphate (pH 7.2), 150 mM NaCl, 2 M Urea, 1 mM TCEP, 20 mM imidazole, and disrupted by sonication. The cell lysates were clarified via centrifugation at 40,000 × g for 50 min and supernatants were applied to $Ni^{2+}$-NTA resin and the column washed with buffer C and eluted with buffer C supplemented with 300 mM imidazole. The eluted proteins were dialyzed against buffer C without imidazole for 3 h to remove imidazole. Following dialysis, Ulp1 enzyme was added to the protein solution for 1 h to remove the SUMO tag and 200 mM BME was added for 16 h to achieve GryA fusion hydrolysis. The solution was passed over $Ni^{2+}$-NTA resin to capture cleaved SUMO and GryA proteins. Flow through containing Piece 3 was collected, supplemented with 1% TFA and purified by HPLC (C4 preparative column). Fractions were analyzed by RP-HPLC and intact ESI-MS and pure fractions were lyophilized.

### Preparation of SEA resin

bis(2-sulfanylethyl)amido (SEA) polystyrene resin was made following the protocol from Ollivier et al, 2014.

**Piece 2 TDP-43 peptides (Strategy 1 residues 315-328; Strategy 2 residues 328-340)**

***Strategy 1, Piece 2 peptides***

Sequence of TDP-43 WT (315-328, A315Thz)- SEA: Thz-FSINPAMMAAAQA-SEA
Sequence of TDP-43 F316meF (315-328, A315Thz, F316meF)- SEA: Thz-meFSINPAMMAAAQA-SEA
Sequence of TDP-43 S317meS (315-328, A315Thz, S317MeS)- SEA: Thz-FmeSINPAMMAAAQA-SEA
Sequence of TDP-43 I318meI (315-328, A315Thz, I318MeI)- SEA: Thz-FSmeINPAMMAAAQA-SEA
Sequence of TDP-43 N319meN (315-328, A315Thz, N319meN)- SEA: Thz-FSImeNPAMMAAAQA-SEA
Sequence of TDP-43 A321meA (315-328, A315Thz, A321meA)- SEA: Thz-FSINPmeAMMAAAQA-SEA
Sequence of TDP-43 M322meM (315-328, A315Thz, M322meM)- SEA: Thz-FSINPAmeMMAAAQA-SEA
Sequence of TDP-43 M323meM (315-328, A315Thz, M323meM)- SEA: Thz-FSINPAMmeMAAAQA-SEA
Sequence of TDP-43 A324meA (315-328, A315Thz, A324meA)- SEA: Thz-FSINPAMMmeAAAQA-SEA
Sequence of TDP-43 A325meA (315-328, A315Thz, A325meA)- SEA: Thz-FSINPAMMAmeAAQA-SEA
Sequence of TDP-43 A326meA (315-328, A315Thz, A326meA)- SEA: Thz-FSINPAMMAAmeAQA-SEA
Sequence of TDP-43 Q327meQ (315-328, A315Thz, Q327meQ)- SEA: Thz-FSINPAMMAAAmeQA-SEA
Sequence of TDP-43 A328meA (315-328, A315Thz, A328meA)- SEA: Thz-FSINPAMMAAAQmeA-SEA

***Strategy 2, Piece 2 peptides***

Sequence of TDP-43 WT (328-340, A328Thz)- SEA: Thz-ALQSSWGMMGML - SEA
Sequence of TDP-43 A329meA (328-340, A329meA, A328Thz)-SEA Thz-meALQSSWGMMGML – SEA
Sequence of TDP-43 L330meL (328-340, L330meL, A328Thz)-SEA Thz-AmeLQSSWGMMGML – SEA
Sequence of TDP-43 Q331meQ (328-340, Q331meQ, A328Thz)-SEA Thz-ALmeQSSWGMMGML – SEA
Sequence of TDP-43 S332meS (328-340, S332meS, A328Thz)-SEA Thz-ALQmeSSWGMMGML – SEA
Sequence of TDP-43 S333meS (328-340, S333meS, A328Thz)-SEA Thz-ALQSmeSWGMMGML – SEA
Sequence of TDP-43 W334meW (328-340, W334meW, A328Thz)-SEA Thz-ALQSSmeWGMMGML – SEA
Sequence of TDP-43 G335meG (328-340, G335meG, A328Thz)-SEA Thz-ALQSSWmeGMMGML – SEA
Sequence of TDP-43 M336meM (328-340, M336meM, A328Thz)-SEA Thz-ALQSSWGmeMMGML – SEA

Sequence of TDP-43 M337meM (328-340, M337meM, A328Thz)-SEA Thz-ALQSSWGMmeMGML – SEA
Sequence of TDP-43 G338meG (328-340, G338meG, A328Thz)-SEA Thz-ALQSSWGMMmeGML – SEA
Sequence of TDP-43 M339meM (328-340, M339meM, A328Thz)-SEA Thz-ALQSSWGMMGmeML – SEA

where, SEA = bis(2-sulfanylethyl)amido group, me = $N^{\alpha}$-methyl, Es = ester, Thz = thiazolidine

**Peptide synthesis**

All fluorenylmethyloxycarbonyl (Fmoc)-protected amino acids were purchased from Oakwood Chemical or Combi-Blocks. For peptide synthesis, bis(2-sulfanylethyl)amino polystyrene resin was prepared in house using trityl chloride resin (Chem-Impex). All analytical reversed-phase HPLC (RP-HPLC) was performed on an Agilent 1260 series instrument equipped with a quaternary pump and an XBridge Protein BEH C4 column (3.5 µm, 4.6 × 150 mm; Waters) at a flow rate of 1 mL/min. Similarly, semi preparative scale purifications were performed employing a XBridge Peptide BEH C4 OBD semi-preparative column (5 µm, 10 mm × 250 mm, Waters) at a flow rate of 4 mL/min. Preparative RP-HPLC was performed on an Agilent 1260 series instrument equipped with a preparatory pump and a XBridge Peptide C4 OBD preparatory column (10 µM; 19 × 250 mm, Waters) at a flow rate of 20 mL/min. All instruments were equipped with a variable wavelength UV-detector. All RP–HPLC steps were performed using 0.1% (trifluoroacetic acid, TFA, Oakwood Chemical) in $H_2O$ (Solvent A) and 90% acetonitrile (Sigma-Aldrich), 0.1% TFA in $H_2O$ (Solvent B) as mobile phases. For LC/MS analysis, 0.1% formic acid (Sigma-Aldrich) was substituted for TFA in mobile phases. Mass analysis was carried out for each product on an LC/MSD (Agilent Technologies) equipped with a 300SB-C18 column (3.5 µM; 4.6 × 100 mm, Agilent Technologies) or a X500B QTOF (Sciex).

All peptides containing C-terminal SEA were synthesized via solid-phase peptide synthesis on a CEM Discover Microwave Peptide Synthesizer (Matthews, NC) using the Fmoc-protection strategy on SEA resin (0.16 mmol/g; Iris Biotech). SEA resin was washed and swelled with N,N dimethylformamide (DMF, Oakwood Chemical) and bubbled in $N_2$ for 15 min respectively. For manual loading of the first amino acid to SEA resin, Fmoc-alanine (10 eq; for Strategy 1 peptides) or Fmoc-leucine (10 eq; for Strategy 2 peptides), HATU (10eq, Oakwood Chemical), and DIPEA (30eq, Sigma-Aldrich) were mixed in DMF and the resin was bubbled with $N_2$ for 1 h. This step was repeated with fresh reagents to ensure complete loading. The resin was then washed with DMF and bubbled in acetic anhydride:DIPEA (20 eq:40 eq) in DMF for 20 min to quench unreacted sites. Subsequent peptide synthesis reactions were performed on an automated, microwave synthesizer. For coupling reactions, amino acids (5 eq) were activated with DIC (5 eq, Oakwood Chemical):Oxyma (5 eq, Oakwood Chemical) and heated to 90 °C for 2 min while bubbling with nitrogen gas in DMF. Fmoc deprotection was carried out with 20% piperidine (Sigma-Aldrich) in DMF supplemented with 0.1 M HOBt (Oakwood Chemical) at 90 °C for 1 minute while bubbling with nitrogen gas. Resin cleavage was performed with 95% TFA, 2.5% triisopropylsilane (TIS, Sigma-Aldrich), and 2.5% $H_2O$ for 3 h at 25 °C. The crude peptide was then precipitated by the addition of a 10-fold volume of ice-cold ether and centrifuged at 4,000 RCF for 10 min at 4 °C. To oxidize the C-terminal SEA, the pellet was washed with ice-cold ether and resuspended with buffer containing 6 M guanidine-HCl, 0.1 M sodium phosphate and

5% (v/v) DMSO at pH 7.5-8.0. This reaction was nutated at 25 °C for 16 h and SEA ring oxidation was confirmed by a -2 Da mass change via LC-MS. To deprotect Thz ring, 200 mM of O-methylhydroxylamine HCl (Combi-Blocks) was added to the reaction mixture and the pH of the reaction was adjusted to 4.0 and incubated at 37 °C for 1 h. Deprotection of the thiazolidine ring was confirmed by a -12 Da mass change via LC-MS. Oxidized and Thz deprotected peptides were purified via preparative C4 RP-HPLC. Fractions were analyzed on analytical C4 RP-HPLC and ESI-MS and those containing pure product (>95%) were pooled, lyophilized, and stored at -80 °C.

**Assembly of semi-synthetic TDP-43 constructs**

To scan $N^{\alpha}$-methyl amino acids from amino acids 316-328, we assembled all Strategy 1 pieces described above. To scan $N^{\alpha}$-methyl amino acids from amino acids 329-339, we assembled all Strategy 2 pieces described above. Two separate strategies were required as we were unable to synthesize a single peptide from residues 316-339 on the requisite SEA resin at sufficient yields. The following three piece assembly strategy was carried out for Strategy 1 and Strategy 2 assemblies.

Piece 1 (bearing MES thioester) and Piece 2 (bearing free N-terminal cysteine and oxidized SEA ring) were combined at ~10 mM Piece 1 and ~8 mM Piece 2 in a degassed buffer of 6 M guanidine-HCl, 0.1 M sodium phosphate, and 100 mM 2,2,2-trifluoroethanethiol (TFET, Sigma-Aldrich) and adjusted to pH 7.0. Excess Piece 2 was used to push all peptide to ligated product and ensure efficient separation of ligated product from starting material during purification. Reactions were incubated at 37 °C for 4 h and progress was monitored via C4 RP-HPLC and ESI-MS analysis. Note the C-terminal oxidized SEA ring remained intact and inert throughout this reaction. Ligation products were purified on a semi-preparative C4 RP-HPLC column and fractions containing target mass were pooled, lyophilized, and stored at -80 °C. This product is referred to as 'Piece 1+2'.

The lyophilized Piece 1+2 construct was resuspended in 0.3-1 ml of degassed buffer (final Piece 1+2 concentration ~0.5 mM) containing 6 M guanidine-HCl, 0.1 M sodium phosphate, 200 mM MES-Na (Sigma-Aldrich), and 50 mM TCEP and adjusted to pH 4.0. This reaction was incubated at 37 °C for 16 h to convert the C-terminal SEA moiety to a MES-thioester as confirmed by a -5 Da mass shift via LC-MS. Once MES conversion was complete, Piece 3 (bearing an N-terminal cysteine) was directly added to this reaction mixture to a concentration of ~1.5 mM. Excess Piece 3 was used to push all Piece 1+2 to ligated product and ensure efficient separation of ligated product from starting material during purification. Reactions were supplemented with 20 mM TCEP and 100 mM TFET, adjusted to pH 7.0, and incubated 37 °C for 16 h. Upon completion of the ligation reaction as judged by RP-HPLC and ESI-MS analysis, excess TFET was removed by dialyzing the reaction mixture against a buffer containing 6 M guanidine-HCl and 0.1 M sodium phosphate at pH 4.0 for 3 h. Desulfurization of the final product to the native TDP-43 LCD sequence achieved through free radical-mediated desulfurization by addition of 200 mM TCEP, 60 mM VA-044 radical initiator and 150 mM of reduced glutathione. Reactions were adjusted to pH 7.0 and incubated at 37 °C for 16 h. Ligation products were purified on a semi-preparative C4 RP-HPLC column and pure fractions were pooled, lyophilized, and stored at -80 °C. Final purities of >95% were judged by analytical RP-HPLC, ESI-MS analysis and SDS-PAGE.

**TDP-43 LCD phase-separated droplet formation**

All His-TDP-43 LCD wild type constructs, mutation variants, and $N^{\alpha}$-methyl variants were dissolved in a buffer containing 25 mM Tris-HCl (pH 7.5), 150 mM NaCl, 6 M guanidine-HCl, 10 mM β-mercaptoethanol (β-ME) and diluted to 300 μM with same buffer. For glycine scanning mutagenesis experiments, droplet formation was induced by diluting all constructs 30-fold in a buffer containing 25 mM Tris-HCl (pH 7.5), 150 mM NaCl, and 10 mM β-mercaptoethanol (β-ME) in the presence of 0 M, 0.4 M, 0.8 M or 1.2 M urea. Solutions were loaded onto a clear bottom Corning Costa 384-well plate and imaged on a ZOE Fluorescent Cell Imager (Bio-Rad).

For $N^{\alpha}$-methyl scanning experiments, droplet formation was induced by diluting all constructs 30-fold in a buffer containing 25 mM Tris-HCl (pH 7.5), 150 mM NaCl, 10 mM β-ME, and 5 μM $NiCl_2$ in the presence of 0 M, 0.4 M, 0.8 M or 1.2 M urea. The formed droplets were immediately loaded onto a clear bottom 384-well plate and absorbance at 600 nm was measured to determine turbidity. Triplicate measurements were performed for all constructs at each condition. Following turbidity measurements, droplets were also imaged on a ZOE Fluorescent Cell Imager.

**Transient transfection of U2OS cells**

U2OS cells were cultured in a Dulbecco's modified Eagle's medium (DMEM) supplemented with 10% fetal bovine serum (FBS). The U2OS cells were seeded on a 35 mm glass bottom confocal dish one day prior to transfection. Cells were transfected with FLAG-GFP-TDP-43 plasmid (or mutations thereof) via lipofectamine 3000 (ThermoFisher). After 24 h, cells were washed twice with PBS, incubated with Hoechst staining reagent (diluted in PBS) for 10 min and imaged by confocal microscopy. Cells were also harvested in SDS sample loading buffer for western blotting (α-FLAG-HRP; MilliporeSigma) to confirm similar expression levels of each TDP-43 variant.

**Creation of TDP-43 inducible expression cell line and cell viability assay**

All constructs were transfected into a Tet inducible cell line, U2OS-TR. The U2OS-TR cells were grown in Tet negative FBS to minimize leaky expression. Cells ($1 \times 10^6$) were plated 24 h prior to transfection. DNA (1 μg) was transfected with Lipofectamine 3000 (ThermoFisher) according to manufacturer's protocol. After 24 h, cells were split into 10 x 100 mm plates in media containing G418 at a concentration of 700 μg/mL. Selection continued for approximately 3 weeks (refreshing G418 twice per week). Single colonies were picked using clonal rings, propagated, and doxycycline induced expression levels were analyzed. For FLAG-GFP-TDP-43 induction, doxycycline was added to a concentration of 1 μg/mL for 24 h. Clonal populations were harvested in SDS sample loading buffer for western blotting (α-FLAG) to confirm similar expression levels of each TDP-43 variant. Colonies from each variant that exhibited similar FLAG-GFP-TDP-43 expression levels were used in the viability assays.

Each stable expression cell line was plated in quadruplet into 7 × 96-well plates at 4,000 cells/well. Induction proceeded as previously mentioned using doxycycline starting at 24 h after plating. Plates were assayed for cell viability via CellTiter-Glo (Promega) according to manufacturer’s protocol (CellTiter-Glo / Promega) each day for 7 days.

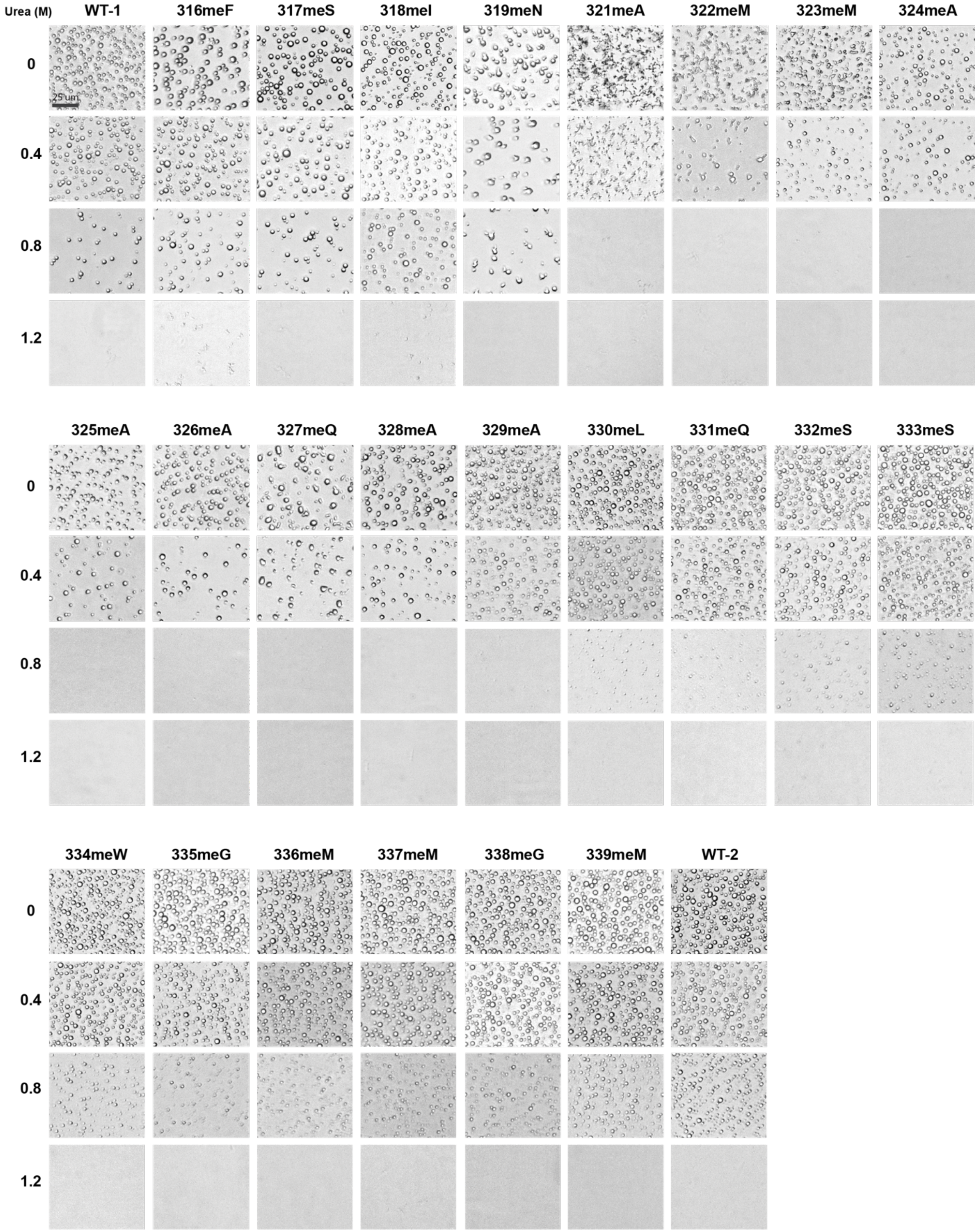


**Fig. S1. Assays of phase separation of TDP-43 variants bearing $N^{\alpha}$-methyl amino acid variants at individual sites.**

Semi-synthetic proteins corresponding to the TDP-43 LC domain were assembled according to the legend to Fig. 1. Individual variants contained a single, m ethyl-capped peptide backbone nitrogen atom. Following purification, samples were incubated under conditions of neutral pH and physiological levels of monovalent salt suitable for protein self-association and phase separation. In addition to normal buffer, each protein sample was also tested in buffer supplemented with 0.4 M urea, 0.8 M urea or 1.2 M urea. Scale bar = 25 µm.

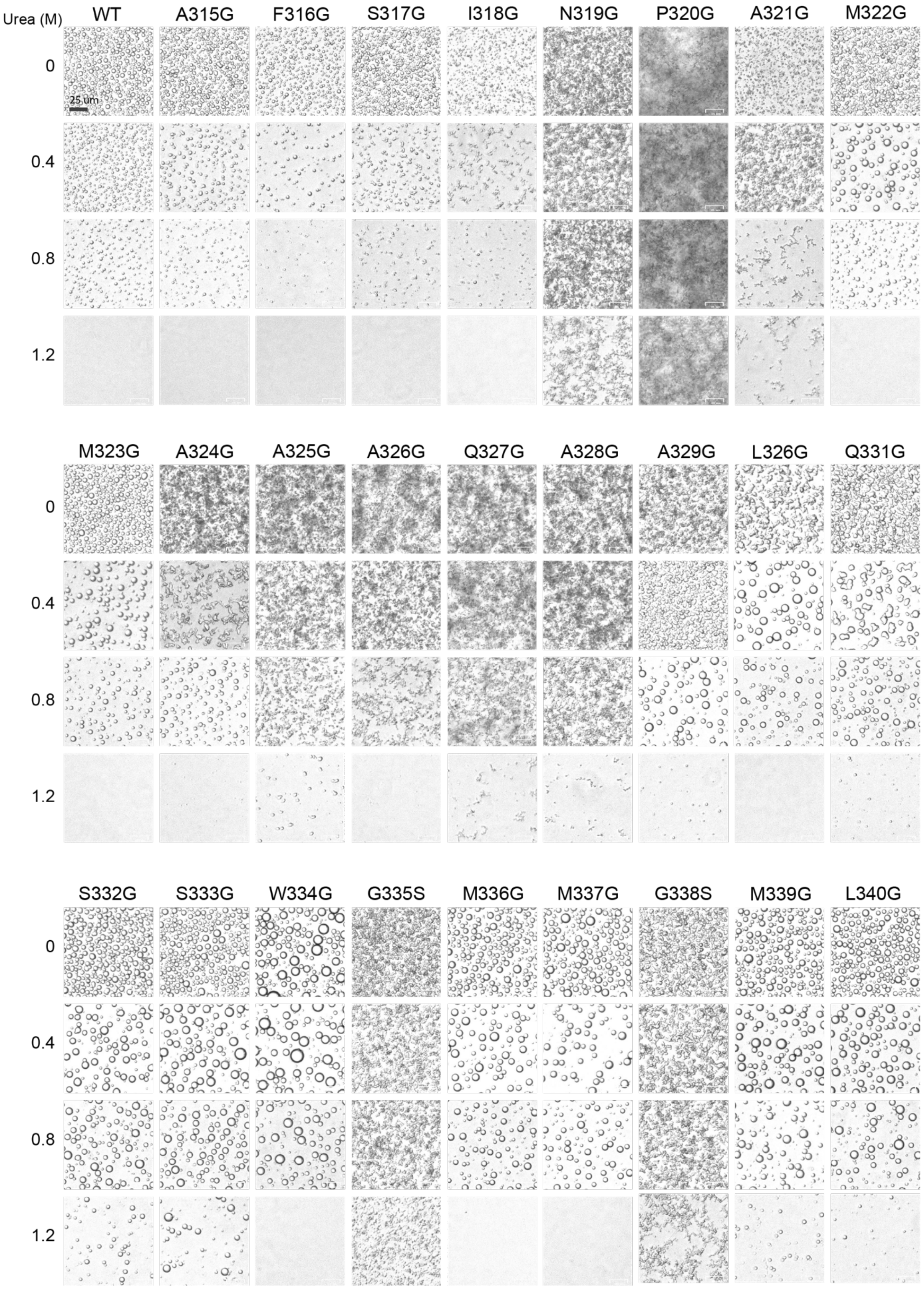

Urea (M)
WT
A315G
F316G
S317G
I318G
N319G
P320G
A321G
M322G
0
0.4
0.8
1.2
25 um
M323G
A324G
A325G
A326G
Q327G
A328G
A329G
L326G
Q331G
S332G
S333G
W334G
G335S
M336G
M337G
G338S
M339G
L340G

**Fig. S2. Assays of phase separation of glycine scanning variants across the ultra-conserved region of the TDP-43 LC domain.**

Protein variants replacing individual amino acid residues of the native TDP-43 LC domain with glycine were expressed in bacterial cells, purified and incubated under conditions of neutral pH and physiological levels of monovalent salt suitable for protein self-association and phase separation. The two glycine residues native to the TDP-43 LC domain, G335 and G338, were changed to serine. In addition to normal buffer, each protein sample was also tested in buffer supplemented with 0.4 M urea, 0.8 M urea or 1.2 M urea. Scale bar = 25 μm.

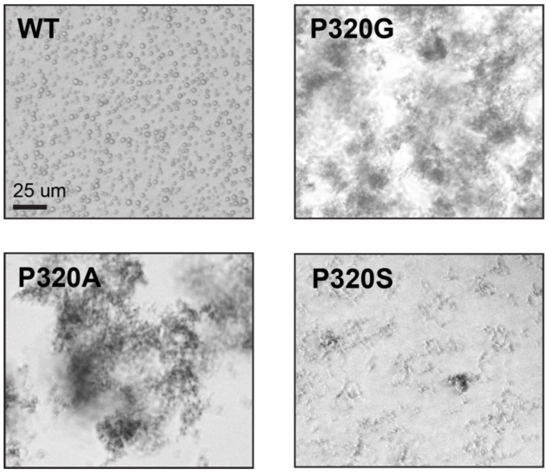


**Fig. S3. Effects of mutational change of proline residue 320 of the TDP-43 low complexity domain on phase separation.**

The native (WT) TDP-43 LC domain and variants changing proline residue 320 to glycine, alanine or serine were assayed for the formation of phase separated liquid-like droplets in aqueous buffer of neutral pH supplemented with 150 mM NaCl. The native protein formed spherical, liquid-like droplets. The three variants changing proline 320 to glycine, alanine or serine formed tangled precipitates. Scale bar = 25 µm.
